# Characterization of novel *Chrysosporium morrisgordonii* sp. nov., from bat white-nose syndrome (WNS) affected mines, northeastern United States

**DOI:** 10.1101/591503

**Authors:** Tao Zhang, Ping Ren, XiaoJiang Li, Sudha Chaturvedi, Vishnu Chaturvedi

## Abstract

Psychrotolerant hyphomycetes including unrecognized taxon are commonly found in bat hibernation sites in Upstate New York. During a mycobiome survey, a new fungal species, *Chrysosporium morrisgordonii* sp. nov., was isolated from bat White-nose syndrome (WNS) afflicted Graphite mine in Warren County, New York. This taxon was distinguished by its ability to grow at low temperature spectra from 6°C to 25°C. Conidia were tuberculate and thick-walled, globose to subglobose, unicellular, 3.5-4.6 µm ×3.5-4.6 µm, sessile or borne on narrow stalks. The stalks occurred at right angles to the hypha. Intercalary aleuroconidia were also observed; the hyphae were septate, hyaline, smooth-walled, 1-3 µm wide, straight or branched. The ITS1-5.8S-ITS2 sequence of *C. morrisgordonii* differed from the closest match to *C. speluncarum* by 37 nucleotides (94% identity); the Neighbor-joining phylogenetic tree confirmed that *C. morrisgordonii* was placed in a well-supported monophyletic group with *C. speluncarum*. LSU and ITS phylogenetic trees further indicated that *C. morrisgordonii, C. speluncarum, Renispora flavissima, Neogymnomyces virgineus and Gymnoascus demonbreunii* formed a distinct group in the *Onygenales.*

## Introduction

During a mycobiome survey from bat’s White-nose syndrome (WNS) affected caves and mines in Upstate New York [1], we isolated an interesting psychrotolerant hyphomycete, which belonged to the genus *Chrysosporium*. This isolate was recovered from soil and sediments collected from Graphite mine in Warren County, New York, USA.

The genus *Chrysosporium*, in the subphylum Pezizomycotina, comprises an extensive collection of cosmopolitan saprotrophs, which are commonly recovered from soil, marine sediments, decaying wood, skin or feet of reptiles [2-5]. Corda first introduced the genus *Chrysosporium* in 1833. Currently, a total of seventy species are recognized r[3]. *Chrysosporium* species are often keratinolytic, and occasionally cause skin and nail infections in humans and animals [2, 6-10]. The resemblance of *Chrysosporium* species to dermatophytic fungi, and their pathogenic potential is of high diagnostic importance [11-15]. The genus is readily identified by mostly white to yellowish colonies, typical conidiophores structures with terminal or intercalary conidia [16]. The conidia are usually subglobose, pyriform, or clavate and are released rhexolytically [2, 16].

In the current study, we describe the new species *Chrysosporium morrisgordonii* based upon the colony, microscopic, physiologic, and molecular data in comparison with closely related *Chrysosporium* species.

## Materials and methods

### Sample origin and isolation of the fungus

Site description and sample collection were described previously [1]. Environmental samples including sediment or swab sample were performed according to the method described previously [1]. Culture and purification process was described in detail previously [1].

### Fungal strains and media preparation

A total of five strains from different reference culture collections were included in the study. Among them, three including *Neogymnomyces demonbreunii* (Ajello et Cheng) CBS 122.67 (ATCC^®^18028™), *Renispora flavissima* (Sigler *et al*.,) CBS 708.79 (ATCC^®^38504™), and *Amauroascus volatilis-patellus* (Eidam) Schroeter CBS 249.72 (ATCC^®^18898™) were provided by the Centraalbureau voor Schimmelcultures (CBS) Utrecht, Netherlands. *Chrysosporium speluncarum* CCF 3760 and *Neogymnomyces virgineus* JN038186 were generous gifts from Prof. Alena Novakova (Institute of Microbiology, Czech Republic) and Prof. Sabrina Sarrocco (Università di Pisa, Italy), respectively. The strains were cultured on potato dextrose agar (PDA), Sabouraud dextrose agar (SDA; peptone 10 g, dextrose 40 g, agar 20 g L^-1^), and malt extract agar (MEA; malt extract 20 g, peptone 1.0 g, dextrose 20 g, agar 20 g L^-1^), respectively. The basal medium (1 L volume) constituents included (NH_4_)_2_SO_4_ 3.0 g, K_2_HPO_4_ 1.0 g, KH_2_PO_4_ 1.0 g, MgSO_4_.7H_2_O 0.5 g, 10 mL trace elements solution [17]. All strains were generally incubated at 20°C or 30°C for 14 days.

### Morphological identification

The growth rate of the isolate was determined at 6°C, 10°C, 15°C, 20°C, 25°C, 30°C on SDA and MEA. The Petri dishes were inoculated in the center, and the colony diameters (in millimeters) were measured daily. The strains were identified using standard methods based on morphological characteristics [2]. The morphological structures were measured in warmed lactophenol mounts. Photomicrographs were obtained with an Olympus Provis AX70 microscope. Scanning electron microscopy techniques were described previously by Figueras and Guarro [18]. The pure culture is deposited in the Mycology Laboratory Culture Collection Repository, Wadsworth Center, New York State Department of Health, Albany, New York, USA.

### DNA extraction and PCR amplification

Genomic DNA from fungal mycelium grown on SDA plates was extracted according to the manufacturer’s instructions (Epicentre, Madison, WI, USA, Cat. No. MC85200). Complete ITS1-5.8S-ITS2 (ITS) and partial 28S rDNA were amplified using universal primers. ITS1 and ITS4 [19] were used to amplify ITS, and NL1 and NL4 [20] were used for partial 28S (large subunit, LSU) amplification. Each PCR reaction mixture (30 µL) contained 3.0 µL 10×PCR buffer, 0.9 µL sense primer (10 µM), 0.9 µL antisense primer (10 µM), 1.0 µL dNTP (10 mM each), 2.0 µL DNA template, 0.3 µL Q5^®^ High-Fidelity DNA Polymerase (2.0 U/ µL) (New England Biolabs, Inc.) and 21.9 µL sterile water. The PCR reactions were performed in an C1000 Touch™ Thermal Cycler (Bio-Rad) with cycling conditions of an initial activation at 98°C for 30 sec, followed by 28 cycles of denaturation at 98°C for 10 sec, annealing at 55°C (ITS) and 58°C (28S) for 30 s, and elongation at 72°C for 25 sec, with a final extension step at 72°C for 2 min. PCR products were purified with the Qiaquick^®^ PCR Purification Kit (Qiagen) and sequenced with the ABI BigDye terminator chemistry in an ABI 3730 DNA sequencer (Applied Genomic Technologies Core, Wadsworth Center). The same primers used in PCR were employed as sequencing primers. The PCR amplicons were assembled, edited using Sequencher 4.6 software (Gene Codes Corp., Ann Arbor, MI, USA) and BLAST searched against two databases, i.e., GenBank (www.ncbi.nlm.nih.gov/) and CBS-KNAW (www.cbs.knaw.nl/).

### Phylogenetic analysis

All sequences were aligned with Clustal X version 1.8 [21] and BioEdit v7.0.9 [22], and the multiple alignments were adjusted manually following visual inspection and the areas of sequence ambiguity were eliminated when necessary. Databases of ITS and 28S were combined for sequence analyses. Phylogenetic analyses using the maximum parsimony analysis (MP) was performed with the MEGA 5.1 computer program [23]. The robustness of branches was assessed by bootstrap analysis with 1,000 replicates. The Dictionary of the Fungi and UniProt (http://www.uniprot.org/taxonomy/) served as the source of taxonomic references for fungal species [24].

### Physiological study

Growth on dermatophyte test medium agar (DTM) and color changes from yellow (acidic) to red (basic) were recorded. Lipase activity was tested by growing on Tween 80 opacity test medium (TOTM) according to Slifkin [25]. Production of urease was determined on Christensen’s urea agar after incubation for 7 d [26]. Amylase, chitinase, proteinase, cellulase, and β-glucosidase activity was verified by growing strains on induction media including basal medium with 0.5% starch, 0.5% chitin, 0.5% casein, 0.5% CM-cellulose, and 0.5% D-(+)-cellobiose, respectively. Cycloheximide tolerance was evaluated by growing strains on SDA supplemented with 0.2% and 0.5% of cycloheximide (Sigma, USA). Tolerance of NaCl was evaluated by the growth of strains on SDA, amended with 3% and 5% NaCl. Hemolysis was evaluated by culturing strains on blood agar. All detections were performed at 20°C for 14 d. Temperature sensitivity tests are incubated at 6°C and 30°C, respectively.

### Nucleotide Sequence Accession Numbers

All nucleotide sequences of the test isolate were deposited in GenBank under accession numbers: ITS sequence (accession number: KM273791); for 28S rRNA gene partial sequence (accession number: KM273792).

## Results

### Morphology

Colonies were moderately fast growing, attaining a diameter of 38-42 mm on MEA, and 32-36 mm on PDA medium after 20-day. Optimal growth was observed at 20°C. At 25°C colonies reached 10-14 mm on MEA and 9-12 mm on PDA in 20 days. Although no growth was observed at 30°C, initial growth was detected at 6°C after 14 days of incubation with colonies attaining diameters of 17-22 mm after four weeks. Colonies on MEA at 20 °C were white to yellowish white, powdery and dense, with a thin margin, slightly cottony at center, with reverse pale yellow to brown (Figure 1A, B). On PDA medium, colonies were yellow to sand colored, powdery, suede-like or slightly granular, raised in the center, with a poorly defined margin with reverse brown to grey in age (Figure 1C, D).

**Fig. 1.**
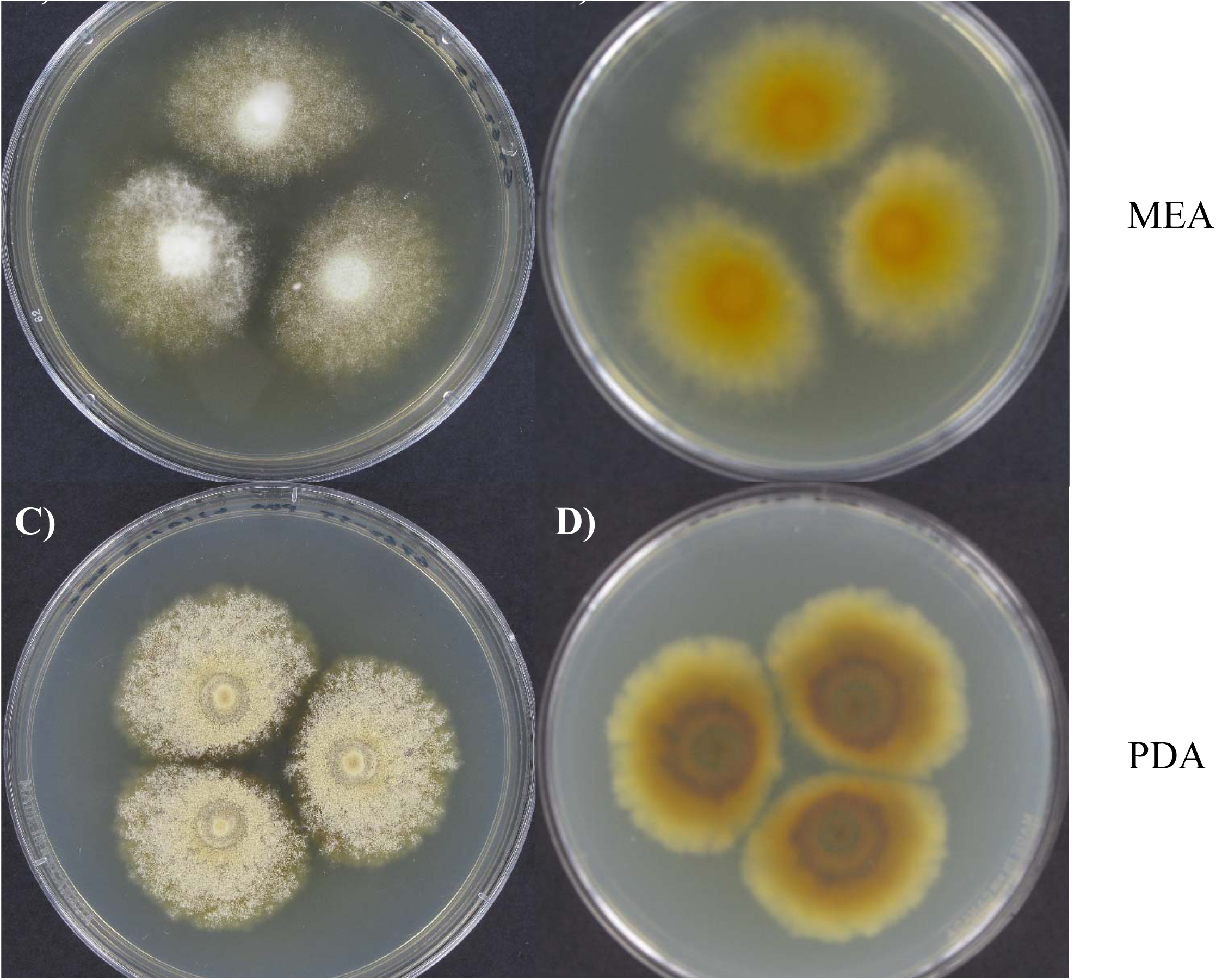
*Chrysosporium morrisgordonii* Zhang, Ren, Li, Chaturvedi, Chaturvedi, sp. nov. A, B) Colony on MEA agar (surface and reverse); C, D) Colony and PDA agar (surface and reverse).

Microscopically, the isolate was distinguishable. Conidia were tuberculate and thick-walled, globose to subglobose, unicellular, 3.5-4.6 µm ×3.5-4.6 µm, sessile or borne on narrow stalks (Figure 2A, 2B). The stalks occurred at right angles to the hypha. Intercalary aleuroconidia were also observed; the hyphae were septate, hyaline, smooth-walled, 1-3 µm wide, straight or branched (Figure 2C-2G).

**Fig. 2.**
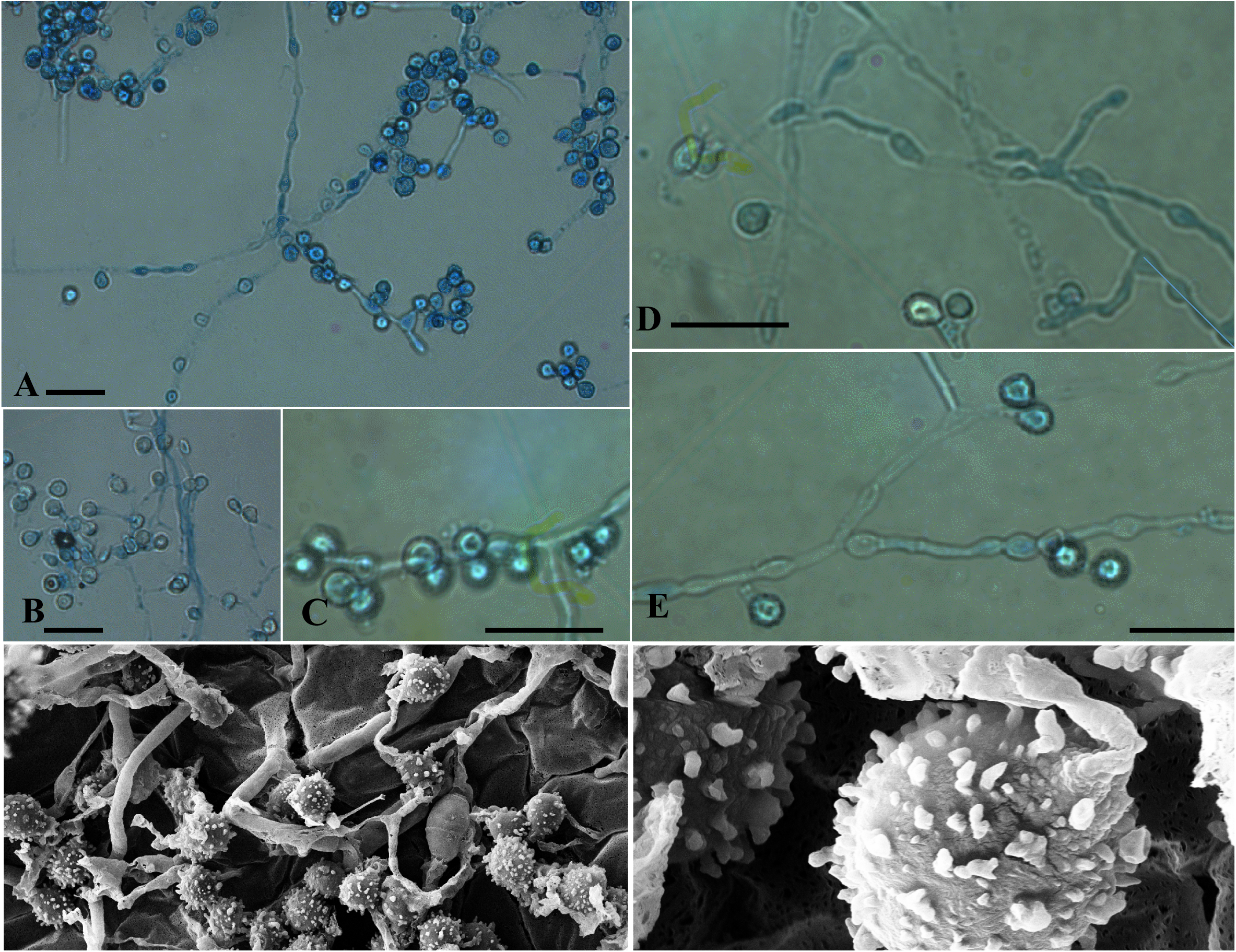
*Chrysosporium morrisgordonii* Zhang, Ren, Li, Chaturvedi, Chaturvedi, sp. nov. A-D) Fertile hyphae bearing sessile conidia; E) Two intercalary aleuroconidia and a terminal chain of tuberculate conidia (white arrows show the septa); F) sessile (some 2-celled) conidia. — Scale bars = 10 μm (A–F, differential interference contrast) or 1 μm (G).

### Phylogeny

The ITS1-5.8S-ITS2 region sequenced from the fungal isolate included 557 base pairs. A search on GenBank database using BLAST revealed only insignificant hits from GenBank, the closest match to an identified *Chrysosporium* strain was *Chrysosporium speluncarum* CCF 3760 with 94% identity (accession no. AM949568), followed by *Gymnoascus demonbreunii* CBS 122.67 (accession no. AJ315842, 84% coverage/84% identity), *Amauroascus volatilis-patellus* CBS 249.72 (accession no. AJ390378, 73% coverage/85% identity), and *Renispora flavissima* CBS 708.79 (accession no. AF299348, 61% coverage/86% identity). None of which were considered significant for a conspecific isolate (>97% identity). The sequence of the strain *C. morrisgordonii* differed from the sequence of *C. speluncarum* CCF 3760 at 37 nucleotide positions. The ITS phylogenetic tree inferred from the nucleotide analysis showed *C. morrisgordonii* grouped in a distinct clade with *C. speluncarum* CCF 3760 (Figure 3).

**Fig. 3.**
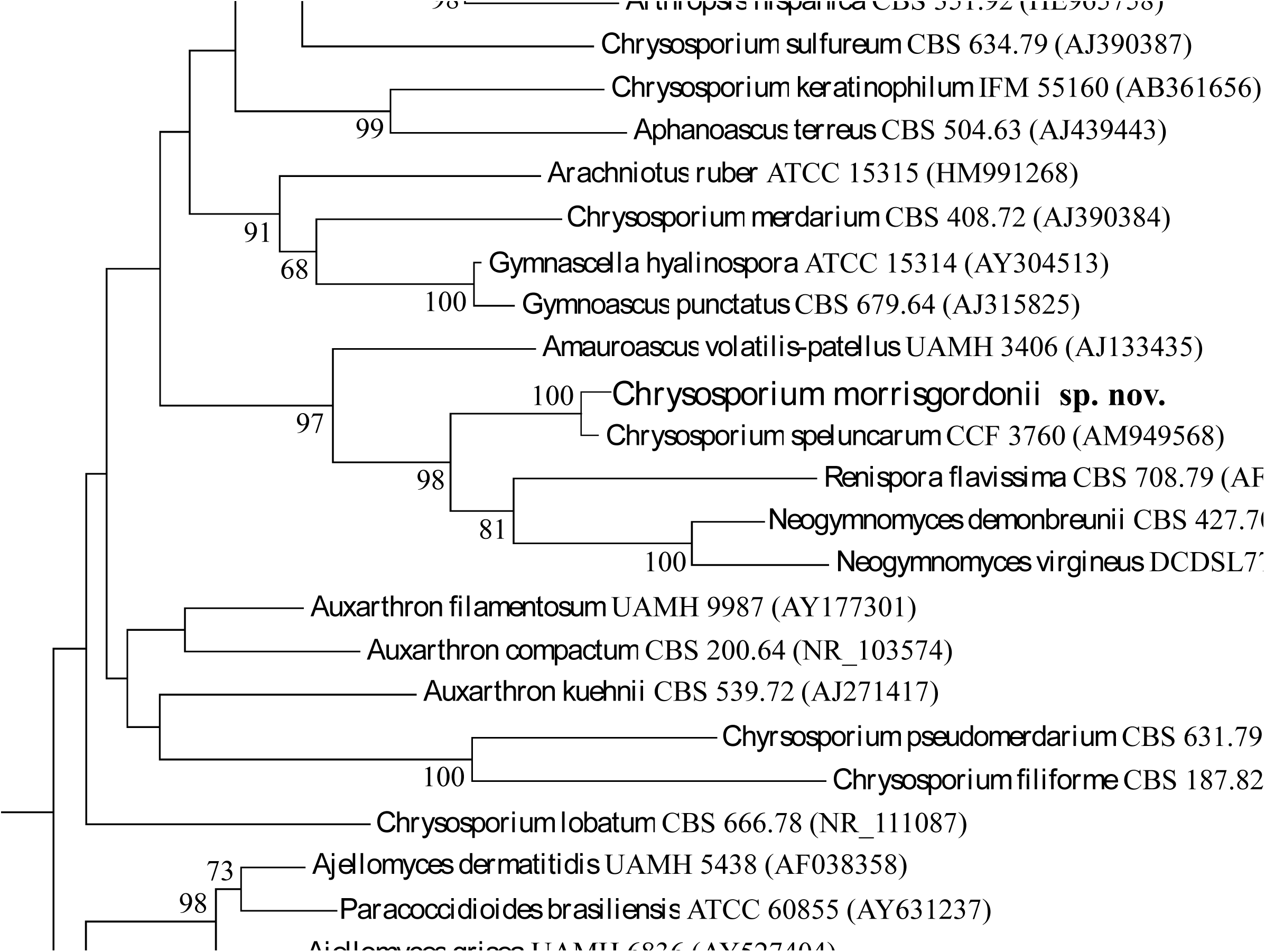
Neighbor-joining tree based on Kimura 2-p corrected nucleotide distances among ITS1-5.8S-ITS2 rRNA gene sequences of the species related with *Chrysosporium morrisgordonii*. Branch lengths are proportional to distance. Bootstrap replication frequencies over 60% (1,000 replications) are indicated at the nodes. The sequences were retrieved from CBS (www.cbs.knaw.nl/) or GenBank (www.ncbi.nlm.nih.gov). *Aspergillus niveoglaucus* CBS 101750 and *Aspergillus alliaceus* NRRL 20602 were used as outgroup.

The D1/D2 region (partial sequence) also did not return a significant BLAST match. The closest sequence in GenBank was *Chrysosporium speluncarum* CCF 3760 (accession no. AM949568, 94% coverage/96% identity) sequence using BLAST search, followed by *Neogymnomyces virgineus* JN038186 (accession no. JN038186), *Amauroascus volatilis-patellus* CBS 249.72 (accession no. AB075324), *Gymnoascus demonbreunii* CBS 122.67 (accession no. AY176716) all displaying 95% similarity. The phylogenetic tree inferred from the analysis of the D1/D2 sequences showed a similar topology, confirming that the isolate formed a well-supported monophyletic group with *Chrysosporium speluncarum* CCF 3760 as the closest neighbor (Figure 4).

**Fig. 4.**
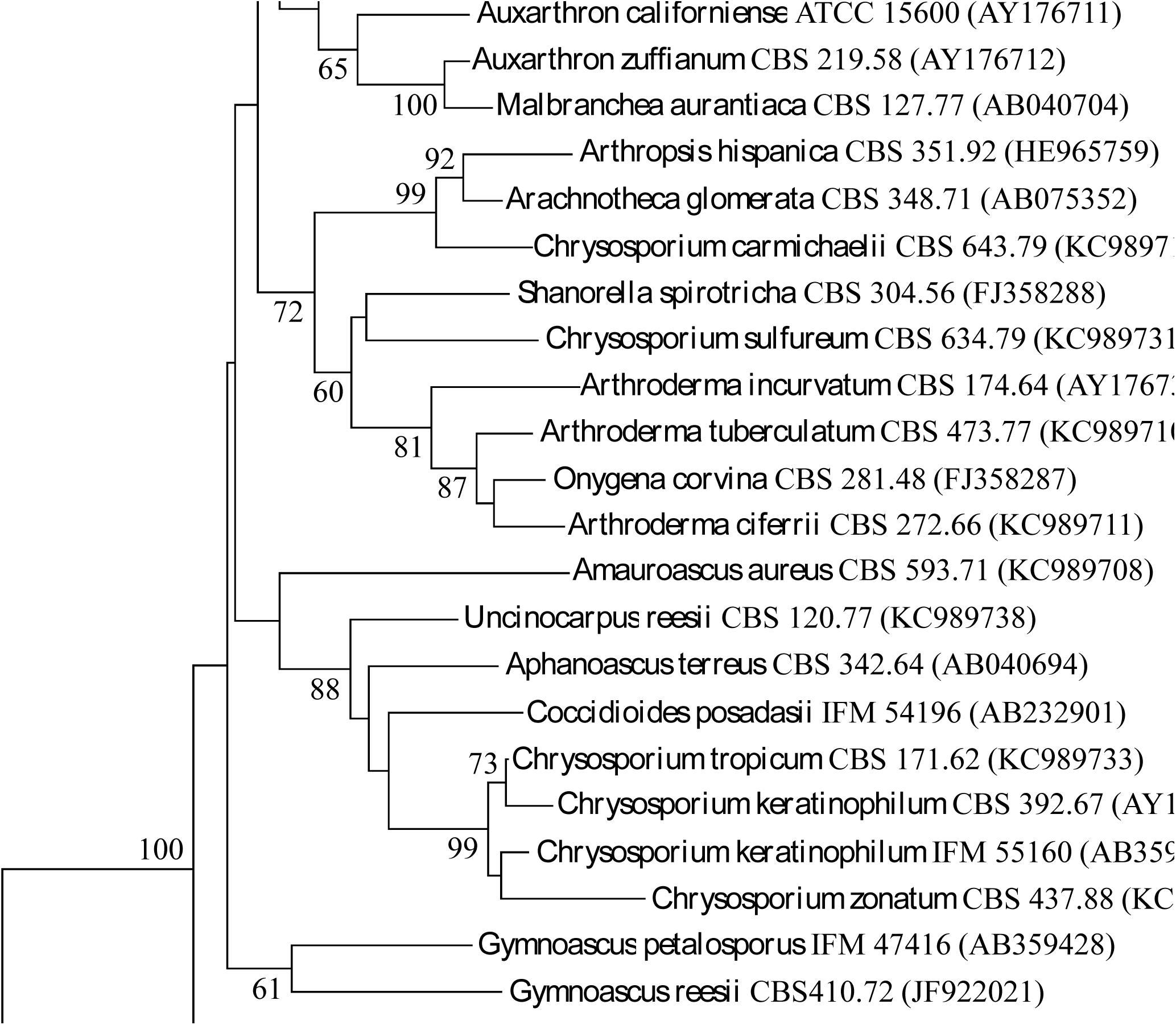
Neighbor-joining tree based on Kimura 2-p corrected nucleotide distances among the D1 and D2 domains of the 28S rRNA gene sequences of the species related with *Chrysosporium morrisgordonii*. Branch lengths are proportional to distance. Bootstrap replication frequencies over 60% (1,000 replications) are indicated at the nodes. The sequences were retrieved from CBS (www.cbs.knaw.nl/) or GenBank (www.ncbi.nlm.nih.gov). *Aspergillus acristata* NRRL 2394 and *Aspergillus fumigatus* CBS 286.95 were used as outgroup.

### Physiology

A summary of crucial physiological tests performed on *C. morrisgordonii* and comparator fungi are presented in Table 1.e All the strains in the study produced hemolysis on blood agar, showed lipolytic, proteolytic, amylolytic, cellulolytic, and chitinolytic activity. *Chrysosporium morrisgordonii* and other fungi grew on casein agar except for *C. speluncarum*. All the fungi tested grew on DTM changing the color of the medium from yellow to red (data not shown), while the strain *C. morrisgordonii* displayed weak activity. The urease and glucosidase tests were positive for all the isolates. On SDA medium with 3% and 10% NaCl, growth was scarce for *A. volatilis-patellus* and *R. flavissima*, while the other four isolates showed good growth (Table 1). Cycloheximide tolerance and different temperature tolerance are listed in Table 1, *Chrysosporium morrisgordonii* and *C. speluncarum* CCF 3760 were psychrotolerant strains, which showed good growth at 6°C, restricted growth at 25°C, and no growth at 30°C.

**Table 1.** Key physiological features of fungi included in this study

| Fungi<br>Physiology | *Chrysosporium*<br>*morrisgordonii* | *C. speluncarum*<br>CCF 3760 | *G. demonbreunii*<br>CBS 427.70 | *N. virgineus*<br>DCDSL7716 | *A. volatilis-patellus*<br>CBS 249.72 | *R. flavissima*<br>CBS 708.79 |
| --- | --- | --- | --- | --- | --- | --- |
| casein agar | ++ | - | ++ | ++ | +++ | + |
| BHI agar | +++ | +++ | +++ | +++ | ++ | +++ |
| lipase | ++ | + | ++ | ++ | +++ | + |
| ureae | +++ | +++ | ++ | +++ | +++ | + |
| chitinase | ++ | ++ | + | +++ | +++ | + |
| cellulase | ++ | +++ | +++ | ++ | ++ | + |
| β-Glucosidase | ++ | + | ++ | ++ | + | + |
| DTM agar | R/B, + | R/B, +++ | R/B, ++ | R/B, ++ | R/B, +++ | R/B, ++ |
| SDA+3% NaCl | +++ | +++ | +++ | +++ | + | + |
| SDA+10% NaCl | + | + | + | + | - | - |
| SDA+0.2% Cyc | +++ | ++ | ++ | +++ | ++ | ++ |
| SDA+0.5% Cyc | +++ | ++ | + | ++ | + | ++ |
| SDA at 6°C | ++ | ++ | - | - | - | - |
| SDA at 30°C | - | - | +++ | +++ | +++ | ++ |

Note: *C*, *Chrysosporium*; *N*, *Neogymnomycetes*; *G*, *Gymnoascus*; *A*, *Amauroascus*; *R*, *Renispora*. BHI agar, Brain Heart Infusion agar; Cyc, cycloheximide; SDA, sabouraud dextrose agar; +++, positive or strong activity; +, scarce positive, moderate activity; -, absence or no activity. Y/A, yellow (acidic); R/B, red (basic).

Phylogenetic analysis showed that *C. morrisgordonii* is closely related to *C. speluncarum* CCF 3760, *G. demonbreunii* CBS 122.67, *A. volatilis-patellus* CBS 249.72, *R. flavissima* CBS 708.79, and *N. virgineus* JN038186. *C. morrisgordoniican* is easily distinguished from *A. volatilis-patellus, R. flavissima*, and *N. virgineus* by growth at 6°C and no growth at 30°C. *Chrysosporium speluncarum* produced diffusible pigment and a larger colony than *C. morrisgordonii* at 25°C while the latter grew slightly better than *C. speluncarum* at 6-10°C (Figure 5).

**Fig. 5.**
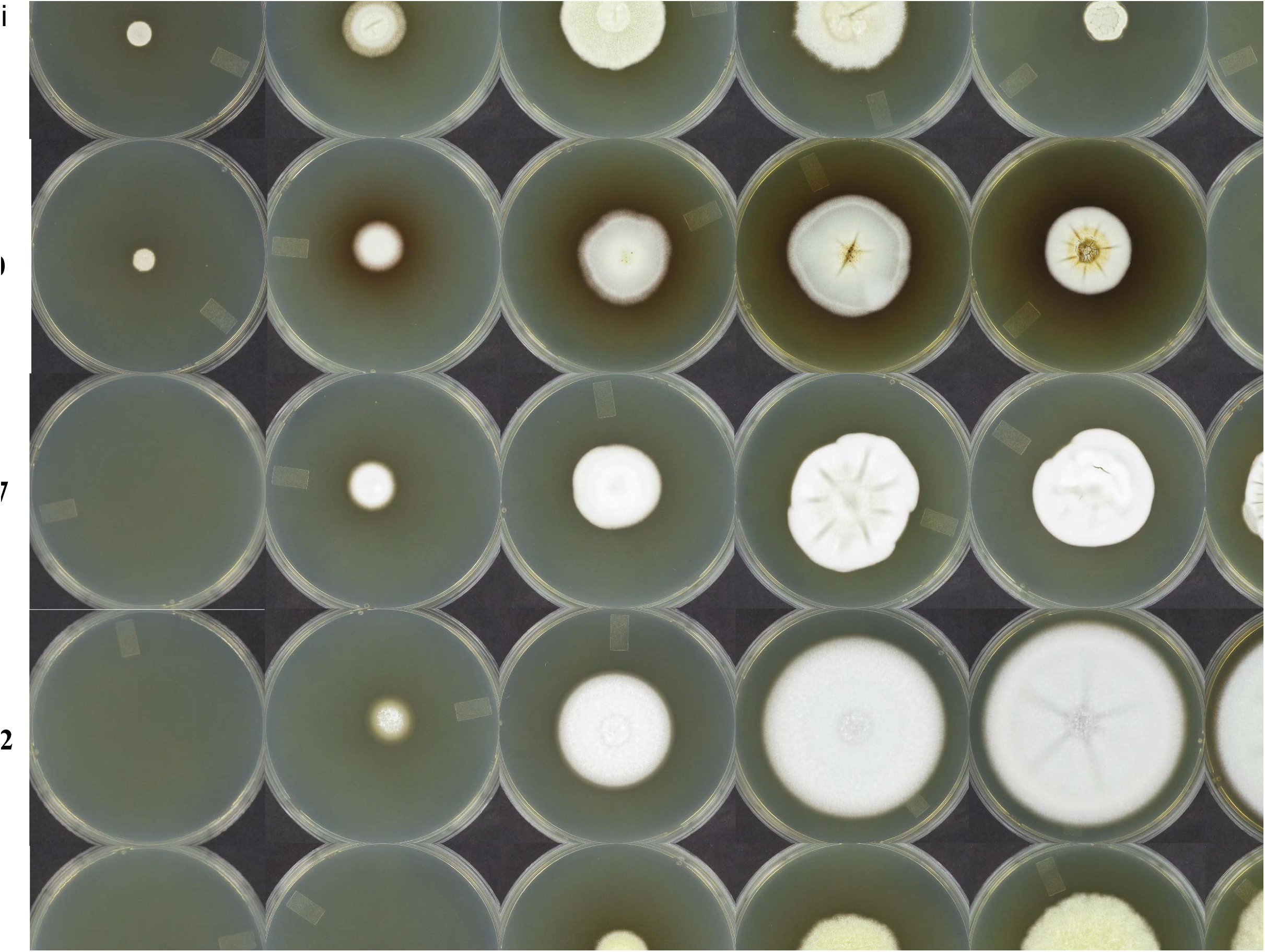
Comparison of the radial growth of C. morrisgordonii at temperatures ranging from 6°C to 30°C on Sabouraud dextrose agar (SAB). Also included are *C. speluncarum* CCF 3760, *G. demonbreunii* CBS 122.67, *A. volatilis-patellus* CBS 249.72, *R. flavissima* CBS 708.79, and *N. virgineus* DCDSL7716.

### Taxonomy

The combined morphological, molecular and physiological features of the strain isolated from the Graphite mine soil samples do not correspond to any of the species described to date within the genus *Chrysosporium* or closely related genera. Thus, we propose here as a new species.

### *Chrysosporium morrisgordonii* Zhang, Ren, Li, Chaturvedi, Chaturvedi, sp. nov. MycoBank MB 830215

#### Etymology

The epithet is derived from late Dr. Morris A. Gordon, eminent mycologist, and ex-director of Mycology Laboratory, New York State Department of Health, where the isolate *Chrysosporium morrisgordonii* was characterized.

#### Holotypus

USA, New York, Warren County, soil,and sediment from Graphite mine, January 2011, ex Morrisgordonii, Isotypus 6762-S5.

*Colonies* on MEA at 20°C attaining a diameter of 35-42 mm after 7 d, whitish to yellowish white, powdery and dense, with rich aerial mycelium, elevated at the center and radially folded, compact; the reverse was pale yellow to brown. *Hyphae* hyaline, septate, smooth-walled, straight or branched, 1-3 µm in diameter. *Conidia* unicellular, tuberculate and thick-walled, globose to subglobose, 3.5-4.6 µm ×3.5-4.6 µm, sessile or borne on at the ends of narrow stalks, 3.5-4.6 µm ×3.5-4.6 µm; aleuriconidia (intercalary conidia) common.

*Colonies* on PDA showed features similar to those on MEA, but was more floccose and sand colored, with more sporulation. The optimum growth temperature was 15-20°C, and the minimum was 6°C, with reduced growth at 25°C and no growth at 30°C.

## Discussion

We presented a detailed characterization of a novel species *C. morrisgordonii* from a bat hibernation site.*C. morrisgordonii* is distinguished by its ability to grow at a low-temperature range from 6°C-25°C and by the combined presence of globose or subglobose conidia sessile or borne at the ends of narrow stalks as well as intercalary aleuroconidia. In the LSU and ITS phylogenetic trees, *Chrysosporium morrisgordonii, Chrysosporium speluncarum, Renispora flavissima*, together with two species, *Neogymnomyces virgineus* and *Gymnoascus demonbreunii* formed a distinct group as reported in the previous studies [14, 27-30]. *C. speluncarum* and *R. flavissima* were recovered from cave samples, have an affinity to bat guano, and exhibit a similar phenotype to *C. morrisgordonii* with yellowish colonies and similar tuberculate conidia [14]. The strain *C. morrisgordonii* differs from *C. speluncarum* by the absence of pigment, powdery growth, two different conidia, growth on casein agar, and better growth at 6-10°C (Figure 4). In contrast, *R. flavissima* differentiates from *C. morrisgordonii* for conidial size (6-8×5-8 µm) and better growth at 30°C [31, 32]. *N. virgineus* and *G. demonbreunii* have a broader ecological distribution and a *Chrysosporium* state with smooth conidia [33, 34].

There are few published reports on psychrophilic or psychrotolerant *Chrysosporium* species; *C. magnasporum* and *C. oceanitesii* grew in Antarctic soil [35], *C. evolceanui* in alpine soil [36], and *C. longisporum* from snake [3], but they could be easily differentiated from *C. morrisgordonii* by their distinct morphology.

## Acknowledgments

Special thanks are grateful to Profs. Alena Novakova (Institute of Microbiology, Czech Republic) and Sabrina Sarrocco (Università di Pisa, Italy) for providing the strains *C. speluncarum* CCF 3760 and *N. virgineus* JN038186. We are thankful to the Wadsworth Center Media & Tissue Culture Core for the preparation of various media and reagents and Wadsworth Center Advanced Genomics Core for sequencing of DNA samples. This study was supported financially partly with funds from the National Science Foundation (16-0039-01), as well as National Natural Science Foundation of China (No.31872617), and the Fundamental Research Funds for the Central Universities (No.3332018097).

